# Stereoscopic Rendering in a Head Mounted Display Elicits Higher Functional Connectivity During Virtual Reality

**DOI:** 10.1101/675710

**Authors:** Caroline Garcia Forlim, Lukas Bittner, Fariba Mostajeran, Frank Steinicke, Jürgen Gallinat, Simone Kühn

**Affiliations:** University Medical Center Hamburg-Eppendorf (UKE), Department of Psychiatry and Psychotherapy, Martinistrasse 52, 20246 Hamburg, Germany; University Hamburg, Department of Human-Computer-Interaction, Vogt-Kölln-Str. 30, 22527 Hamburg, Germany; Max Planck Institute for Human Development, Lise-Meitner Group for Environmental Neuroscience, Lentzeallee 94, 14195 Berlin, Germany

**Author notes:** Corresponding author: Simone Kühn.

**Keywords:** virtual reality, stereoscopic and monoscopic googles, fMRI, seed-based functional connectivity, fractional amplitude of low-frequency fluctuations, resting-state networks

## Abstract

Virtual reality (VR) simulates real world scenarios by creating a presence in users. Such immersive scenarios lead to more similar behaviour to that displayed in real world settings, which may facilitate the transfer of knowledge and skills acquired in VR to real world situations. VR has already been used in education, psychotherapy, rehabilitation and it is an appealing choice for training intervention. The aim was to investigate to what extend VR technology can be used in a magnetic resonance imaging scanner(MRI), addressing the question of whether brain connectivity differs between VR and screen via mirror projection presentations. Moreover, we investigate whether stereoscopic goggle stimulation, where eyes receive different input, would elicit more brain connectivity than stimulation where both eyes receive the same input (monoscopic). To our knowledge, there is no previous research addressing this question. Multiple analyses were performed to cover different aspects of brain connectivity: fractional low frequency fluctuation, independent component analysis, seed-based functional connectivity and graph analysis. In goggles (mono and stereoscopic) vs. screen, we found connectivity differences in cerebellum and postcentral gyrus and in visual and frontal inferior cortex in visual/default-mode networks. Considering specific areas, we found higher connectivity between superior frontal cortex and temporal lobe, as well as inferior parietal cortex with calcarine and lingual. Furthermore, superior frontal cortex and insula/putamen were more strongly connected in stereoscopic, in line with our hypothesis. We assume that conditions eliciting most connectivity should be suited for long-term interventions as extended training under these conditions could permanently improve functional/structural connectivity.

## 1 Introduction

Virtual reality (VR) is used in various contexts such as entertainment, education, psychotherapy, rehabilitation and other conditions. In these computer-generated environments the user can perceive, feel and interact in a manner that is similar to a physical place, which is usually achieved by a combination of multiple sensory channels, such as sight, sound and touch. An essential feature of VR is that it creates a sense of presence in its users, which in turn leads to behaviour that is more similar to the behaviour displayed in real world settings. Importantly this feeling of presence may facilitate transfer of knowledge and skills acquired in VR to similar real world situations, which would make VR an ideal choice for training intervention purposes. In preparation for such training studies and to test to what extend VR technology can be used in a magnetic resonance imaging scanner (MRI) we conducted the present study. We set out to investigate whether VR visual stimulation using MRI compatible goggles with 3D stereoscopic stimulation (in which the image is rendered separately for each eye creating the illusion of depth and 3D effect) differs in terms of brain connectivity from more commonly applied presentation forms using goggles with 2D monoscopic presentation (in which both eyes receive the same visual input) and a conventional screen back-projection via a mirror. To our knowledge, there is no previous research using functional MRI to address this question. The previous studies either used different stimulus material to investigate different degrees of spatial presence (Baumgartner et al., 2008; Baumgartner et al., 2006; Dores et al., 2013; Havranek et al., 2012; Lee et al., 2005) and/or used electroencephalography (EEG) (Baumgartner et al., 2006; Dan and Reiner, 2017; Havranek et al., 2012; Kober et al., 2012; Slobounov et al., 2015).

Most interesting for the present endeavour and the question whether VR elicits higher brain connectivity, is an EEG study that compared brain signals during navigation either on a desktop PC (2D) or on a large wall projection in 3D (Kober et al., 2012). The 3D condition was accompanied by higher cortical parietal activation in the alpha band, whereas the 2D condition was accompanied by stronger functional connectivity between frontal and parietal brain regions, indicating enhanced communication. In two additional EEG studies, in which different modes of presentation have been compared, but in which brain activity instead of connectivity was the focus of investigation, contradictory evidence has been gathered. Likewise in a navigation task comparing a condition in which participants wore 3D glasses and watched a screen vs. a 2D screen condition, higher theta power in frontal midline structures was observed in the 3D VR condition (Slobounov et al., 2015). In contrast a study investigating paper folding (origami) learning with 3D glasses vs. with a 2D film showed that the 2D condition displays a higher so-called cognitive load index computed as the ratio of the average power of frontal theta and parietal alpha. A last study focussed on intra-hippocampal EEG recordings comparing real world navigation vs. VR navigation and demonstrated that oscillations typically occurs at a lower frequency in virtual as compared to real world navigation (Bohbot et al., 2017).

Brain regions that have repeatedly received attention in the endeavour to explain differences between VR-related presence experiences in comparison to 2D or less immersive environments are the prefrontal cortex, the parietal cortex as well as the hippocampus (Baumgartner et al., 2008; Baumgartner et al., 2006; Beeli et al., 2008; Bohbot et al., 2017; Dan and Reiner, 2017; Kober et al., 2012). For this reason, we used whole brain connectivity analysis approaches as well as ROI-based approaches focussing on the effects of the type of the display (conventional 2D screen, MRI goggles with monoscopic view effect, MRI googles with stereoscopic view effect) which capture two different degrees of immersion.

## 2 Methods

### 2.1 Participants

The local psychological ethics committee of the University Medical Center Hamburg-Eppendorf, Germany, approved of the study. Twenty-six healthy participants were recruited from a local participant pool. After complete description of the study, the participants’ informed written consent was obtained. Exclusion criteria for all participants were abnormalities in MRI, relevant general medical disorders and neurological diseases. Additional exclusion criteria were movement above the threshold of 0.5 (Power et al., 2012) during the scanning section and completion of all 3 conditions. After the exclusion criteria were applied, the number of subjects dropped to twenty three (mean age = 26.5, SD = 4.8 years, female:male = 11:12).

### 2.2 Task Procedure

While situated in the scanner participants were exposed to a game (FlowVR) that was inspired by the game Flower by Thatgamecompany (http://thatgamecompany.com/flower/) and programmed in Unity (Bittner et al., submitted). The player flies through a nature scene with the goal of making the landscape blossom (Fig. 1). This can be achieved by the player flying close to flowers, which are surrounded by a coloured halo, to virtually pollinate them so that more flowers grow in the surrounding area. Visual and auditory elements associated with positive affect have been implemented. The task has been designed with the goal to reduce negative affect and initiate the experience of flow (Csikszentmihalyi, 2014).

**Figure 1:**
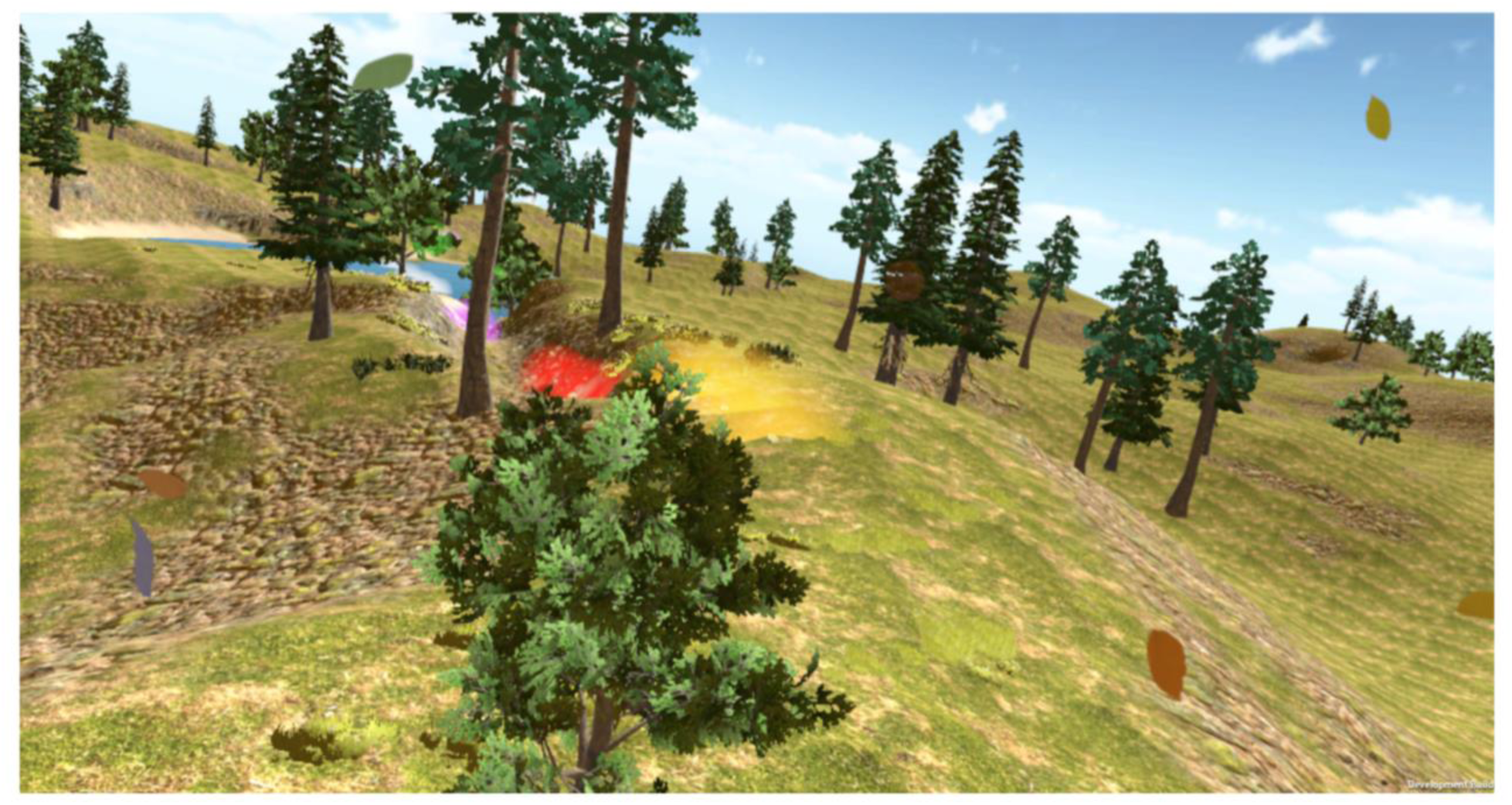
A screenshot of the player’s view when flying in the virtual landscape. The colourful halos indicate the positions of the next flowers to be “pollinated”.

Before entering the scanner, the participants were given information about the gameplay. They were asked to imagine to be a bee, whose goal is to make the landscape flourish by flying above it and touching flowers or trees which are surrounded by a bright halo. That would lead to pollen popping out of the flowers and new flowers, bushes or trees growing all around. They were told to follow the path of the flowers and collect as many flowers as possible, however, they were likewise instructed to not mind when missing out on a flower. Participants used an MR compatible button box with four buttons in a row from Nordic Neuro Lab to navigate in the game. The user had to hold the controller with both hands and use the two left buttons for flying upwards and downwards and the other two for flying to the left and right. The speed of the flight was kept constant. Each run lasted about 5 min.

Participants underwent three conditions, one in which the game was projected from a screen via mirror projection, one in which MRI compatible goggles were used either with a 3D stereoscopic stimulation (in which the image is rendered separately for each eye creating the illusion of depth and 3D effect) and a 2D monoscopic presentation (in which both eyes received the same visual input). MRI compatible googles used were the VisuaStim digital (http://www.mrivideo.com/visuastimdigital.php), with a resolution of 800×600 (SVGA) and filed of view (FoV) of 30 degrees horizontal and 24 degrees vertical. The order of the conditions was randomly assigned to the participants.

### 2.3 Scanning Procedure

Structural images were collected on a Siemens Prisma 3T scanner (Erlangen, Germany) and a standard 32-channel head coil was used. The structural images were obtained using a three-dimensional T1-weighted magnetization prepared gradient-echo sequence (MPRAGE) (repetition time = 2500 ms; echo time = 2.12ms; TI = 1100 ms, acquisition matrix = 240 × 241 × 194, flip angle = 9°; 0.8 × 0.8 × 0.94 mm voxel size). Resting state data was acquired after the T1 image. We acquired whole brain functional images while participants were asked to keep their eyes closed and relax for 5 min. We used a T2*-weighted echo-planar imaging (EPI) sequence (repetition time = 2,000 ms, echo time = 30 ms, image matrix = 64 × 64, field of view = 216 mm, flip angle = 80°, slice thickness = 3.0 mm, distance factor = 20 %, voxel size = 3 × 3 × 3 mm^3^, 36 axial slices, using GRAPPA). Images were aligned to the anterior-posterior commissure line.

### 2.4 Functional MRI Data Analysis

#### 2.4.1 Preprocessing

To ensure for steady-state longitudinal magnetization, the first 10 images were discarded. Slice timing and realignment were performed in the remaining images. The individual anatomical images T1 were coregistered to functional images and segmented into white matter, gray matter, and cerebrospinal fluid. Data was then spatially normalized to Montreal Neurological Institute (MNI) space and spatially smoothed with an 8-mm FWHM to improve signal-to-noise ratio. Signal from white matter, cerebrospinal fluid and movement were regressed. To reduce physiological high-frequency respiratory and cardiac noise and low-frequency drift data was filtered (0.01 – 0.08 Hz) and, finally, detrended. All steps of data preprocessing were done using SPM12 (Wellcome Department of Cognitive Neurology) except filtering that was applied using REST toolbox (Song et al., 2011). In addition, to control for motion, the voxel-specific mean framewise displacement (FD) was calculated according to Power and colleagues (Power et al., 2012). We excluded from the analyses participants who had an FD above the recommended threshold of 0.5.

#### 2.4.2 Fractional Amplitude of Low-Frequency Fluctuation (fALFF)

To investigate voxel-wise changes in the amplitude of low frequency spontaneous fluctuations in the blood-oxygen-level dependent imaging (BOLD) signal in the whole brain, we calculated the fALFF using REST toolbox. Subject-specific fALFF maps were taken to the second level analysis in SPM12.

#### 2.4.3 Independent component analysis (ICA)

We examined the resting-state networks given by the spatial grouping of voxels with temporally coherent activity calculated in a data-driven fashion using ICA. ICA was performed in GIFT software (http://icatb.sourceforge.net/; 24) using Infomax algorithm. The number of spatially independent resting-state networks (N) was estimated by the GIFT software (N=26). The identification of the networks was done automatically using predefined GIFT templates and later the resting-state networks of interested, the default mode network (DMN) and visual networks, were chosen by two specialists (CGF and SK). For every resting-network of interest, subject-specific spatial connectivity maps were taken to the second level analysis in SPM12.

#### 2.4.4 Seed-based Functional Connectivity (SeedFC)

Functional connectivity (FC) is one of the most popular methods to infer connectivity in neuroimaging. When FC is calculated by means of the temporal correlations (Pearson’s correlation) between a region of interest to the other voxels in the whole brain, it is known as seed-based functional connectivity (SeedFC). We investigated the seed-based connectivity maps, using as seed the brain regions of interest in VR, namely, bilateral superior and middle frontal cortex, bilateral hippocampus, bilateral superior and inferior parietal cortex defined in the anatomical automatic labelling (AAL) atlas. Fischer transformation were applied to the individual FC maps obtaining Z scores to improve normality. The Z score maps were taken to the second level in SPM 12.

#### 2.4.5 Graph analysis

To examine differences in the topology of the brain networks we performed graph analysis (Bullmore and Sporns, 2009). The first step was to construct the functional connectivity matrices, where nodes and links should be defined. Nodes were brain regions created based on the AAL116 atlas (Tzourio-Mazoyer et al., 2002) and the links were the connectivity strength between nodes calculated using Pearson’s correlation coefficient. The node-averaged time series used to infer the connectivity strength were extracted for each subject using the REST toolbox (Song et al., 2011). To avoid false positive links, connectivity values that were not statistically significant (p-value > = 0.05) were excluded. Once the functional matrices are built, graph analysis can be applied in order to characterize their topology. At this stage, thresholding was applied, namely density threshold ranging from 0.1 to 0.9 in steps of 0.1. Thresholding means that only links with the highest connectivity strengths are kept until the desired density is reached, e.g. a threshold of 0.1 means 10% of the links with the highest connectivity were kept and the remaining ones were set to 0. Graph analysis were then applied to these thresholded matrices using the Brain Connectivity toolbox (brain-connectivity-toolbox.net). The main graph measures were chosen: betweenness centrality that measures the fraction of all shortest paths that pass through an individual node; characteristic path length which accounts for the average shortest path between all pairs of nodes; efficiency which is the average inverse shortest paths and transitivity that measures the relative number of triangles in the graph, compared to total number of connected triples of nodes. For a complete description of the graph measures please refer to (Bullmore and Sporns, 2009; Rubinov and Sporns, 2010).

#### 2.4.6 Statistics

We were interested in two particular contrasts: (1) VR visual stimulation using MRI compatible goggles with 3D stereoscopic stimulation and 2D monoscopic presentation *vs* screen via mirror projection, to investigate for brain differences during googles and screen, (2) 3D stereoscopic stimulation, in which the image is rendered separately for each eye creating the illusion of depth and 3D effect *vs* 2D monoscopic presentation, most commonly applied presentation in which both eyes receive the same visual input, so that we could investigate the effect of stereoscopic view in the brain.

The resulting maps per subject of each analysis the second level analysis in SPM12 (Statistical Parametric Mapping package; Wellcome Department for Imaging Neuroscience, London, United Kingdom; http://www.fil.ion.ucl.ac.uk/spm) with mean FD as covariate. Using a family wise error (FWE) threshold on the cluster level of *p* < 0.05 we ran the two contrasts in both the positive and the negative direction.

## 3 Results

Since we were mostly interested in the direction of potential brain connectivity differences between conditions, we employed multiple different analysis pipelines. We employed four methods that focus in different aspects of the brain signal, the intrinsic frequency fluctuation and the connectivity given by the spatial grouping of voxels with temporally coherent activity and by the temporal correlations between areas, respectively, fALFF that measures the amplitude of the low frequency fluctuation of the BOLD signal, ICA that uncover the resting-state brain networks, the seed-based FC that calculate the brain network related to specific regions of interest and graph analysis that characterizes the topology of the brain networks. Results are summarized in Table 1.

**Table 1:** Group differences in fALFF, ICA and Seed-FC analyses.

| Analysis | Contrast | Labels | MNI coordinates | $T$ | cluster size (in voxels) | $P$ (cluster level FWE corrected) |
| --- | --- | --- | --- | --- | --- | --- |
| fALFF | googles > screen | Left cerebellum VI | -20 -62 -26 | 4.32 | 92 | 0.007 |
|  |  | Left postcentral | -42 -40 60 | 4.05 | 93 | 0.007 |
|  | googles < screen | Right frontal superior orbital | 18 56 -4 | 6.13 | 72 | 0.006 |
| ICA<br>Primary<br>visual | googles < screen | Left middle occipital | 6 -90 30 | 6.25 | 365 | <<0.001 |
|  | googles > screen | Left calcarine | -8 -94 10 | 4.6 | 152 | 0.018 |
| ICA<br>Higher<br>visual | googles < screen | Bilateral cuneus | 0 -78 24 | 4.7 | 232 | 0.001 |
|  | googles > screen | Left lingual | -4 -64 4 | 6.2 | 257 | <<0.001 |
| ICA<br>DMN | googles > screen | Bilateral lingual | -2 -66 0 | 5.1 | 192 | 0.003 |
|  |  | Inferior frontal gyrus/<br>precentral gyrus | -34 0 28 | 5.0 | 324 | <<0.001 |
| SeedFC<br>Bilateral<br>superior<br>frontal | googles > screen | Left superior temporal<br>pole | -48 16 -12 | 4.7 | 332 | <<0.001 |
|  | stereoscopic > monoscopic | Left insula/putamen | -34 10 10 | 5.4 | 237 | <<0.001 |
| SeedFC<br>Bilateral<br>inferior<br>parietal | googles > screen | Right calcarine | 18 -98 4 | 4.5 | 182 | 0.002 |
|  |  | Right lingual | 26 -88 -6 |  |  |  |
|  |  | Right calcarine | 20 -88 2 |  |  |  |

### 3.1 fALFF

In fALFF we found significantly higher fALFF (Fig. 2) in left cerebellum (VI, −20, −62, −26), left postcentral gyrus (−42, −40 60) in the googles (monoscopic+stereoscopic) condition compared with the screen condition. In the reverse contrast we observed higher fALFF in right superior orbital frontal cortex (−18, 56, −4) (all results family wise error corrected *p*<0.05).

**Figure 2:**
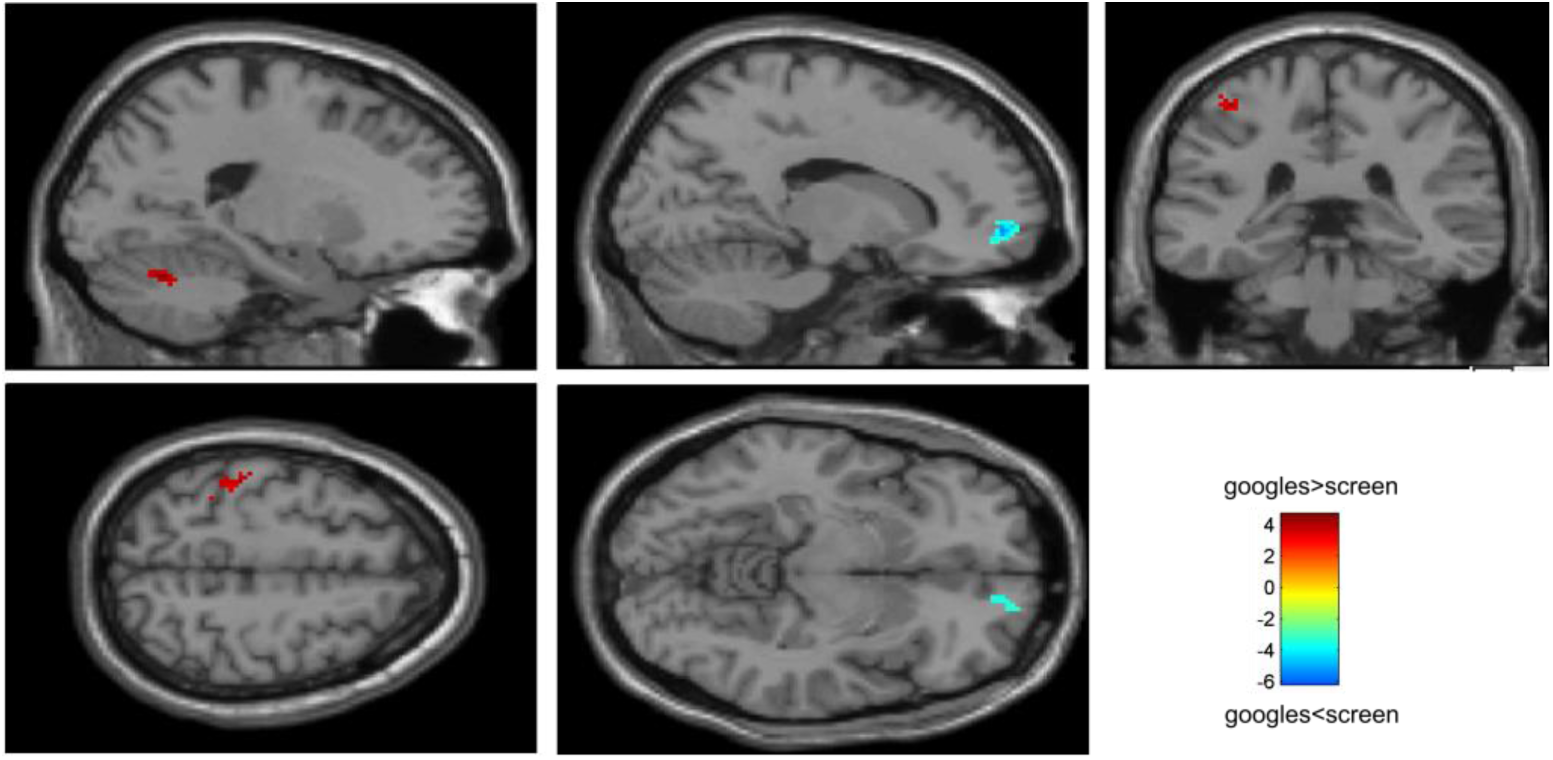
Group differences in fALFF. Higher fALFF (in red) in left cerebellum and left postcentral gyrus in the googles (monoscopic+stereoscopic) condition compared with the screen condition. In the reverse contrast (in blue) there was higher fALFF in right superior orbital frontal cortex.

### 3.2 ICA

In the visual networks (Fig. 3), we found increased in connectivity in the primary and higher networks in the left calcarine (−8, −94, 10) and left lingual (−4, −64, 4), respectively, in the googles (monoscopic+stereoscopic) as compared to screen condition. A decreased in connectivity was observed in the left middle occipital (6, −90, 30) in the primary visual network and in the bilateral cuneus (0, −78, 24) in the higher visual one. Investigating the default mode network (Fig. 3), we found an increase in the connectivity in the inferior frontal (−34, 0, 28) and bilateral lingual (−2, - 66, 0) for googles (monoscopic+stereoscopic) as compared to screen condition. No significant difference between the contrast monoscopic vs stereoscopic was found in the visual networks nor in the DMN. FDR was used to account for multiple comparison correction due to multiple networks.

**Figure 3:**
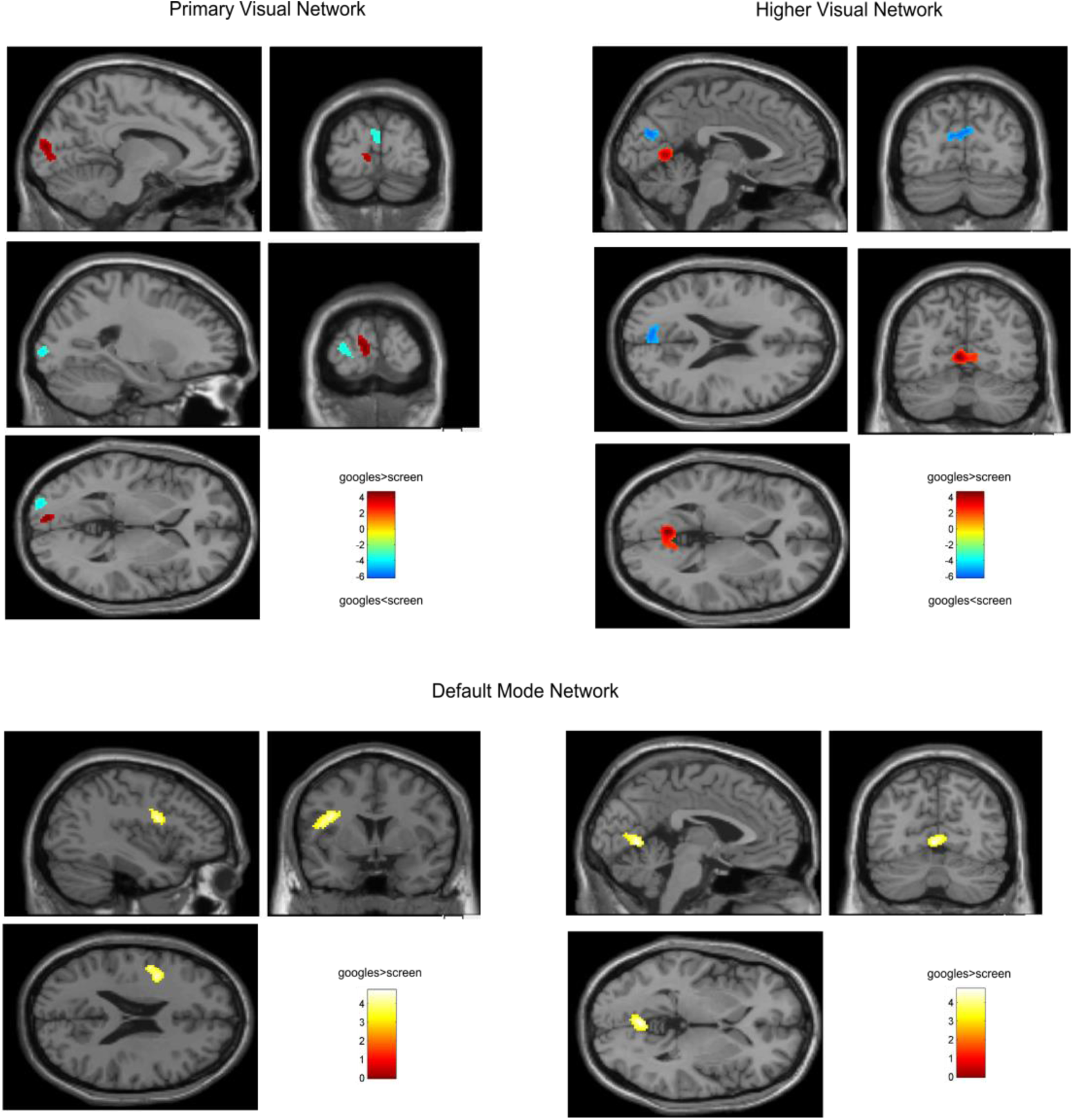
Group differences in the resting state networks: DMN and Visual. In the primary visual network there was an increase in connectivity (in red) in the left calcarine and in the higher visual network in the left lingual in the googles (monoscopic+stereoscopic) condition as compared to the screen condition. A decreased in connectivity (in blue) was seen in the left middle occipital in the primary visual network and in the bilateral cuneus in the higher visual network. In the DMN, we found an increase (in yellow) in the connectivity in the inferior frontal and bilateral lingual for googles (monoscopic+stereoscopic) as compared to screen condition.

### 3.3 SeedFC

In a ROI-based seed analysis we used the following ROIs: bilateral superior frontal cortex, middle frontal cortex, hippocampus, superior parietal cortex, inferior parietal cortex. We found significant seed-based connectivity (Fig. 4) between bilateral superior frontal cortex and the left superior temporal lobe (−48, 16, −12) for the MRI goggle contrast googles (monoscopic+stereoscopic) > screen and to left insula and putamen (−34, 10, 10) in the stereoscopic view contrast (stereoscopic> monoscopic; Fig. 5). The bilateral inferior parietal cortex was more strongly connected to right calcarine cortex (18, −98, 4; 20, −88, 2) and right lingual cortex (26, −88, −6) in the goggles vs screen condition (Fig. 4). All Seed-based ROI analysis results were family wise error corrected at *p*<0.05. To account for multiple comparison correction due to multiple seeds, FDR was used.

**Figure 4:**
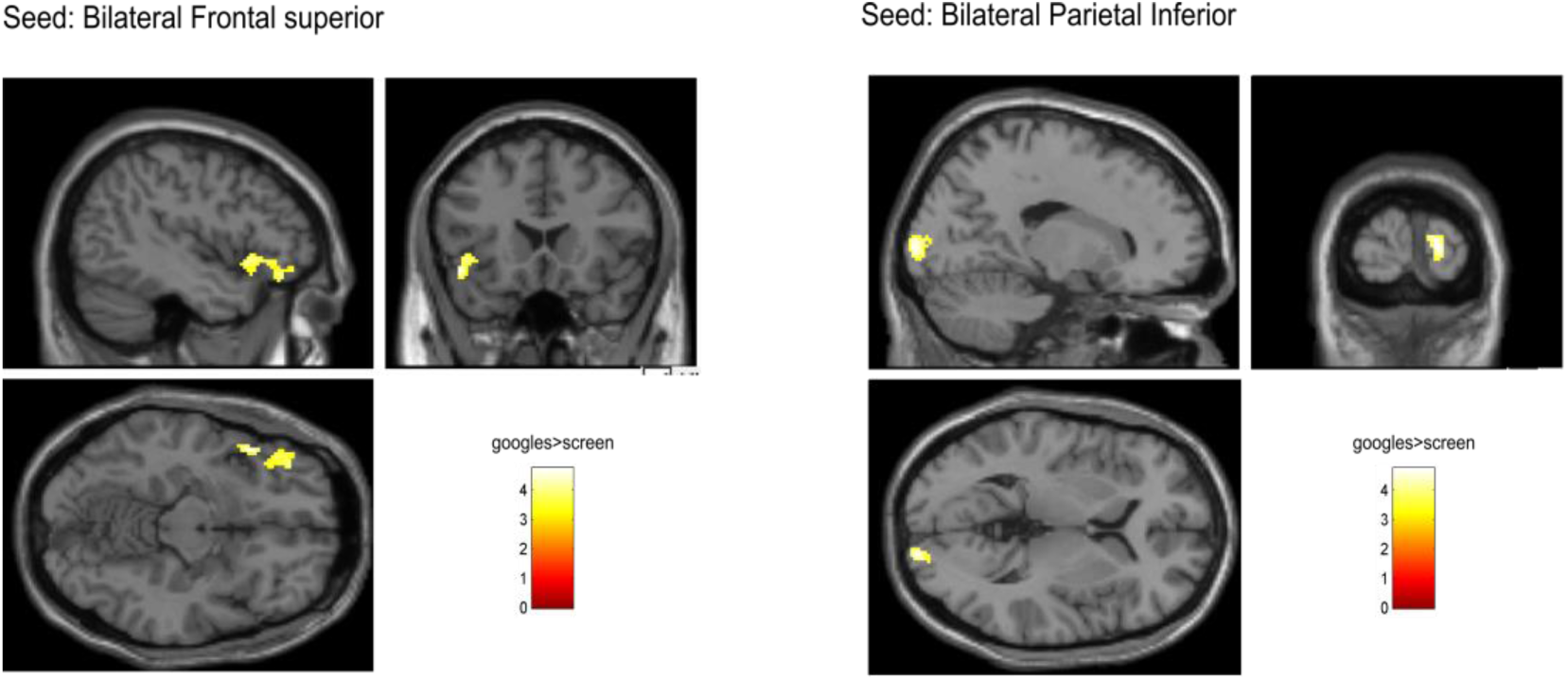
Group differences in SeedFC in googles vs screen condition. There was stronger connectivity between a seed in the bilateral superior frontal cortex and the left superior temporal lobe for the MRI goggle contrast googles (monoscopic+stereoscopic) as compared to the screen. Additionally, a seed in the bilateral inferior parietal cortex was more strongly connected to right calcarine cortex and right lingual cortex in the same contrast.

**Figure 5:**
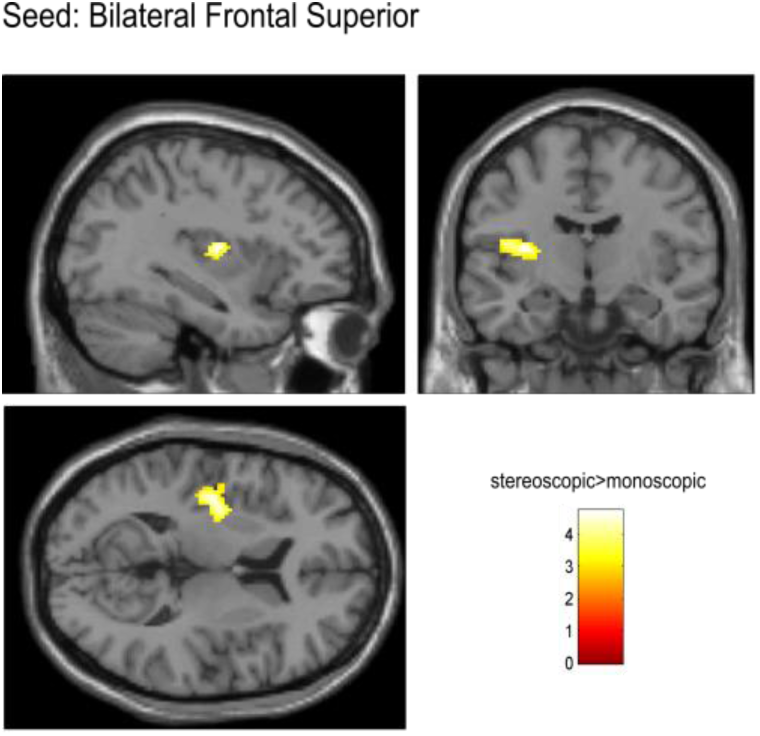
Group differences in the stereoscopic view contrast stereoscopic vs monoscopic in SeedFC for seed in the bilateral superior frontal cortex. The stereoscopic view elicited stronger connectivity between the bilateral frontal cortex and left insula and putamen.

### 3.4 Graph analysis

No topological differences consistent across threshold were found.

## 4 Discussion

Within the scope of the present study we set out to unravel the effects of VR stimulation on functional brain connectivity. In order to do so we used a virtual game in which the player had the task to fly through a natural scene with the goal to make the landscape blossom, which was designed with the goal to decrease negative affect and induce the experience of flow in its players. To disentangle the effects of presenting the visual stimuli via MRI-compatible goggles, we compared the goggle stereoscopic and monoscopic condition with the screen condition. We believe that the goggles, which are mounted on the user’s head and have the ability to display stereoscopic images, are more immersive than a back-projection of a Screen via a mirror system. As in the latter the participant receives only 2D images and can still see the scanner bore and oftentimes even the staff operating the scanner next to the presented stimuli. In order to investigate differences in functional brain connectivity induced by the stereoscopic view, namely the fact that the image is rendered separately for each eye creating the illusion of depth and 3D effect, we contrasted the stereoscopic and monoscopic condition directly. With the goal to obtain a comprehensive picture of brain connectivity we chose four common approaches to analyse resting state fMRI data, namely the assessment of the amplitude of low frequency fluctuation (fALFF, (Zou et al., 2008)), independent component analysis (Calhoun et al., 2001), seed-based functional connectivity analysis and graph analysis (Bullmore and Sporns, 2009).

In line with our hypothesis, the goggles and the stereoscopic contrast revealed stronger brain connectivity for the respective condition with more immersion, that is googles (stereoscopic and monoscopic) compared to screen and stereoscopic compared to monoscopic generally elicited higher brain connectivity. We found higher fALFF in left cerebellum and postcentral gyrus for goggles compared to the screen. In the visual networks, we found an increase in connectivity in the left calcarine and left lingual for the same contrast and in the DMN there was increased connectivity in the inferior frontal cortex and bilateral lingual gyrus. Additionally, in the seed-based analysis we found higher connectivity between bilateral superior frontal cortex and the temporal lobe, as well as bilateral inferior parietal cortex with right calcarine and right lingual cortex for the two goggle vs screen conditions. Furthermore, we found superior frontal cortex and insula/putamen to be more strongly connected in stereoscopic vs monoscopic view. Only from prefrontal cortex we found higher brain activity, that is higher fALFF values in right superior orbital frontal cortex.

These results can be viewed as in line with the hypothesis proposed by Jäncke and colleagues (Jäncke et al., 2009) stating that prefrontal cortex is involved in the experience of presence. In particular bilateral dorsolateral prefrontal cortex (DLPFC) activity was shown to be negatively correlated with the subjective report of the experience of presence (Baumgartner et al., 2008) in adults. Moreover, the authors report the results of an effective connectivity analysis from which they conclude that the right DLPFC is involved in down-regulating the activation in the dorsal visual processing stream. Furthermore the authors interpret the observed increase of activity in the dorsal visual stream during presence experience as a sign of higher action preparation in the virtual reality because the brain responds to it similarly as in real-life situations (Jäncke et al., 2009). However, on the same dataset the left DLPFC was shown to be positively connected to brain regions that are part of the default-mode network (such as medial prefrontal cortex, anterior cingulate cortex, thalamus, brain stem, nucleus caudatus and parahippocampus). Due to the involvement of the latter brain regions in self-referential processing the interpretation is that higher left DLPFC activation when participants experience less presence leads to an up-regulation of self-referential processing which reflects the detachment from the VR experience.

In contrast to this previous study, in which the focus was on the subjective feeling of presence, we set out to investigate differences in brain connectivity between objectively different conditions of stimulation. A major disadvantage of the previous design was that the stimulation used to elicit different degrees of presence was not the same and participants were only passively watching the displays. For this reason, we confronted participants with the same interactive VR game in all three conditions with the difference being the hardware presentation procedures used to display the respective environment.

Our results can be viewed as being in line with the findings and interpretations shown in association to perceived experience of presence (Baumgartner et al., 2008; Jäncke et al., 2009), since we also find more fALFF in right superior orbital frontal cortex in the 2D screen condition compared to the two goggle conditions. Next to these previous fMRI results our results can also be perceived as in line with results from an EEG study in which the same spatial navigation task in a virtual maze was compared between a projection onto a large wall which was supposedly more immersive than a display on a small Desktop PC screen (Kober et al., 2012). The authors report stronger functional connectivity between frontal and parietal brain regions in the Desktop display condition.

However, the rational for the present study was slightly different from the studies presented before. We set out to test whether overall the stereoscopic VR presentation elicits higher degrees of functional brain connectivity than monoscopic and a screen display. Our hypothesis was that the condition that elicits the most brain connectivity should be most suited for long-term brain training interventions, assuming that extended training under these conditions could permanently improve brain connectivity on a functional as well as a structural level. Our results show that the majority of contrasts and functional connectivity indicators resulting from different analysis pipelines reveal higher brain connectivity between brain regions in the goggle condition and the stereoscopic condition in particular, which we interpret as a hint that training in VR environments in contrast to environments displayed on a screen may be superior in eliciting and therewith facilitating brain connectivity in intervention studies.

At present, a major drawback of the implementation of VR in an MRI environment is the fact that the VR experience is limited to the stereoscopic input to the eyes without the experience of movement in space. Usually the stereoscopic view in VR is accompanied by the fact that individuals can freely turn their head and move in space while the visual input is adapted to the individual’s movements. However, since the head cannot be freely moved in the MRI scanner, due to its resulting movement artefacts in the images, the actual differences in brain connectivity between a VR and a screen presentation of an environment might be underestimated. Future research may attempt to use motion tracking systems to enable this movement related visual feedback, while at the same time correct for the occurring motion artefacts in the acquired MRI images (Stucht et al., 2015). The limited field of view and resolution of the MRI compatible goggles introduces yet another limitation to such studies. As it can affect the level of immersion for both monoscopic and stereoscopic conditions. Specifically, to the design of the present study, the question still remains of whether and how frequently the participants noticed the 3D effect when, for example, the bee was not flying close enough to the virtual objects.

## 6 Acknowledgement

The study has been funded by a grant of the German Science Foundation (SFB 936/C7).

SK has been additionally the German Science Foundation (DFG KU 3322/1-1), the European Union (ERC-2016-StG-Self-Control-677804) and a Fellowship from the Jacobs Foundation (JRF 2016-2018).

We thank Karolin Ney for helping in the data acquisition.

